# Learning Peptide Properties with Positive Examples Only

**DOI:** 10.1101/2023.06.01.543289

**Authors:** Mehrad Ansari, Andrew D. White

## Abstract

Deep learning can create accurate predictive models by exploiting existing large-scale experimental data, and guide the design of molecules. However, a major barrier is the requirement of both positive and negative examples in the classical supervised learning frameworks. Notably, most peptide databases come with missing information and low number of observations on negative examples, as such sequences are hard to obtain using high-throughput screening methods. To address this challenge, we solely exploit the limited known positive examples in a semi-supervised setting, and discover peptide sequences that are likely to map to certain antimicrobial properties via positive-unlabeled learning (PU). In particular, we use the two learning strategies of adapting base classifier and reliable negative identification to build deep learning models for inferring solubility, hemolysis, binding against SHP-2, and non-fouling activity of peptides, given their sequence. We evaluate the predictive performance of our PU learning method and show that by only using the positive data, it can achieve competitive performance when compared with the classical positive-negative (PN) classification approach, where there is access to both positive and negative examples.

## 1 Introduction

As short-chain amino acids, peptides have attracted growing attention in pharmaceutics [1–3], therapeutics [4–6], immunology [7–9], and biomaterials design [10–12]. However, the development of novel peptides remains a challenge due to poor pharmacokinetic properties that restrict the design space and necessitate unnatural amino acids or cyclization, increasing the complexity of their design.[13] Computational design and data-driven discovery strategies have arisen as promising low-cost techniques in the pre-experiment phase to expedite the process of generating accurate predictions of peptide properties, and shortlist promising candidates for follow-up experimental validation. Some examples of these successful applications include single nucleotide polymorphisms (SNP) and small-indel calling [14], estimating the impact of non-coding variants on DNA-methylation [15], as well as for the prediction of protein function [16], structure [17, 18], and protein-protein interactions [19]. Sequence-based learning strategies aim at mapping peptide’s natural biological function to its sequence. In a supervised learning setting, this is done by training on sequence-function examples. This means that sequence-function relationships are learned by iteratively training on samples of different classes (i.e. positive and negative examples in binary classification). The performance of the classifier is highly dependent on the quality of the training samples and the ratio of the positive and negative samples [20, 21]. In bioinformatics, a variety of supervised-learning algorithms, such as support vector machines [22], random forest [23], logistic regression [24], and naive Bayesian classifier [25], have been successfully applied to develop classification models.

However, lack of negative examples in numerous biological applications [26–29] limits the feasibility of constructing such reliable classifiers. As an example, medical information records typically contain the positively diagnosed diseases of a patient, and the absence of a diagnostic record does not necessarily rule out a disease for him/her. Most high-throughput screening methods solely focus on identifying the positive examples, thus, it is much more straightforward to confirm a property than to ascertain that it does not hold. As an example, a potential binding site is confirmed if a protein binds to a target, but failure to bind only means that the binding conditions were not satisfied under a given experimental setting. With the technological advances, identifying specific properties can be improved, and biological samples formerly not known to have a property can now be classified with confidence. As an example, Li et al. [30] demonstrated on the changes in protein glycosylation site labeling throughout four time points over 10 years. Another example is protein-protein interaction (PPI) [31, 32], where experimentally validated interacting and non-interacting protein pairs are used as positive and negative examples, respectively. However, the selection of non-interacting protein pairs can be challenging for two reasons: 1. As more novel PPIs are constantly being discovered over time, some non-interacting protein pairs (i.e. negative examples) might be mislabeled. 2. The positive examples are significantly outnumbered by a large number of protein pairs for which no interactions have been identified. Similar situations can be found in drug–drug interaction identification [33], small non-coding RNA detection [34], gene function [35, 36] and phage-bacteria interaction [37] prediction, and biological sequence classification [38, 39].

To address the challenges above, we demonstrate on a positive-unlabeled (PU) learning framework to infer peptide sequence-function relationships, by solely exploiting the limited known positive examples in a semi-supervised setting. Semi-supervised learning techniques are a special instance of weak supervision [40, 41], where the training is based on partially labeled training data (i.e. labeled data can be either positive or both positive and negative samples). PU learning builds classification models by primarily leveraging a small number of labeled positive samples and a huge volume of unlabeled samples (i.e. a mixture of both *positive* (P) and *negative* (N) samples) [42]. Depending on how the *unlabeled* (U) data is handled, existing PU learning strategies are divided into two categories. 1. *Reliable negative identification*: this category identifies *reliable negatives* (RN) within U, and then performs ordinary supervised (PN) learning [43, 44]; 2. *Adapting the base classifier*: this treats the U samples as N with smaller weights (biased learning) and adapts the conventional classifiers to directly learn from P and U samples [45, 46]. The former *reliable negative identification* strategies rely on heuristics to identify the RN, and they have been widely used in none-coding RNA identification [34], none-coding RNA-disease association [47], gene function prediction [35, 48], disease gene identification [26, 49, 50], and single-cell RNA sequencing quality control [51]. On the other hand, *adapting the base classifier* algorithms are Bayesian-based approaches that focus on estimating the ratio of positive and negative samples in U (class prior), which then can be applied for classification using the Bayes’ rule. One major limitation is that their performance largely depends on good choices of weights of U samples, which are computationally expensive to tune [52]. Thus, compared to the first strategy, there has been a fewer use cases of them in the literature [53–55]. An excellent overview of PU leaning strategies can be found in [42]. Li et al. [20] also systematically reviewed the implementation of 29 PU learning methods in a wide range of biological topics.

In this work, we take advantage of the flexibility of reliable negative identification PU strategy, and discover peptide sequences that are likely to map to certain properties. Specifically, we demonstrate on a two-step technique, where Step 1 handles the deficiency of negative training examples by extracting a subset of the U samples that can be confidently labeled as N (i.e. RN). Subsequently, Step 2 involves training a deep neural network classifier using the P and the extracted RN, and applying it to the remaining pool of U. Reliable negative identification in Step 1, is an adaption of the *Spy* technique formerly employed in handling unlabeled text data [43]. In this approach, some randomly selected positive samples are defined as spies, and are intentionally mislabeled as negatives. The reliable negative examples are iteratively found within the unlabeled samples for which the posterior probability is lower than the posterior probability of the spies. We use our approach to predict different peptide properties, such as hemolysis, resistance to non-specific interactions (non-fouling), and solubility.

This manuscript is organized as follows: in Section 2, we describe the datasets, architecture of the deep learning models, and our choices for the hyperparameters. This is followed by evaluating the model in a comparative setting with the classical PN classifier in Section 3. Finally, we conclude the paper in Section. 4, with a discussion of the implications of our findings.

## 2 Materials and Methods

### 2.1 Datasets

#### Hemolysis

Hemolysis is referred to the disruption of erythrocyte membranes that decrease the life span of red blood cells and causes the release of Hemoglobin. It is critical to identify non-hemolytic antimicrobial peptides as a non-toxic and safe measure against bacterial infections. However, distinguishing between hemolytic and non-hemolytic peptides is a challenge, since they primarily exert their activity at the charged surface of the bacterial plasma membrane. In this work, the hemolysis classifier is trained using data from the Database of Antimicrobial Activity and Structure of Peptides (DBAASP v3 [56]). Hemolytic activity is defined by extrapolating a measurement assuming dose response curves to the point at which 50% of red blood cells are lysed. Activities below 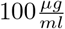, are considered hemolytic. The data contains 9,316 sequences (19.6% positives and 80.4% negatives) of only L- and canonical amino acids. Each measurement is treated independently, so sequences can appear multiple times. This experimental dataset contains noise, and in some observations (*∼* 40%), an identical sequence appears in both negative and positive class. As an example, sequence “RVKRVWPLVIRTVIAGYNLYRAIKKK” is found to be both hemolytic and non-hemolytic in two different lab experiments (i.e. two different training examples).

#### Solubility

This data contains 18,453 sequences (47.6% positives and 52.4% negatives) based on PROSO II [57], where solubility was estimated by retrospective analysis of electronic laboratory notebooks. The notebooks were part of a large effort called the Protein Structure Initiative and consider sequences linearly through the following stages: Selected, Cloned, Expressed, Soluble, Purified, Crystallized, HSQC (heteronuclear single quantum coherence), Structure, and deposited in PDB [58]. The peptides were identified as soluble or insoluble by “Comparing the experimental status at two time points, September 2009 and May 2010, we were able to derive a set of insoluble proteins defined as those which were not soluble in September 2009 and still remained in that state 8 months later.” [57]

#### Non-fouling

Non-fouling is defined as resistance to non-specific interactions, and this data is obtained from [59]. A non-fouling peptide (positive example) is defined using the mechanism proposed in [60]. Briefly, White et al. [60], showed that the exterior surfaces of proteins have a significantly different frequency of amino acids, and this increases in aggregation prone environments, like the cytoplasm. Synthesizing self-assembling peptides that follow this amino acid distribution and coating surfaces with the peptides creates non-fouling surfaces. This pattern was also found inside chaperone proteins, another area where resistance to non-specific interactions is important[61]. Positive data contains 3,600 sequences. Negative examples are based on 13,585 sequences (79.1% of dataset are negatives) coming from insoluble and hemolytic peptides, as well as, the scrambled positives. The scrambled negatives are generated with lengths sampled from the same length range as their respective positive set, and residues sampled from the frequency distribution of the soluble dataset. Samples are weighted to account for the class imbalance caused by the negative examples dataset size. This dataset is gathered based on the mechanism proposed in [60].

#### SHP-2

SHP-2 is a ubiquitous protein tyrosine phosphatase, whose activity is regulated by phosphotyrosine (pY)-containing peptides generated in response to extracellular stimuli. SHP-2 is involved in processes such as cell growth, differentiation, migration, and immune response. [62] The SHP-2 dataset contains fixed-length peptides (5 AA residues) optimized for binding to N-SH2 domain, obtained from [63]. Total dataset size is 300, with 50% positive examples.

### 2.2 Model Architecture

We build a recurrent neural network (RNN) to identify the position-invariant patterns in the peptide sequences, using a sequential model from Keras framework [65] and the TensorFlow deep learning library back-end [66]. In specific, the RNN employs bidirectional Long Short Term Memory (LSTM) networks to capture long-range correlations between the amino acid residues. Compared to the conventional RNNs, LSTM networks with gate control units can learn dependency information between distant residues within peptide sequences more effectively [67–69]. An overview of the RNN architecture is shown in Figure 2. This architecture is identical to the one used in our recent work in edge-computing cheminformatics [64].

**Figure 1:**
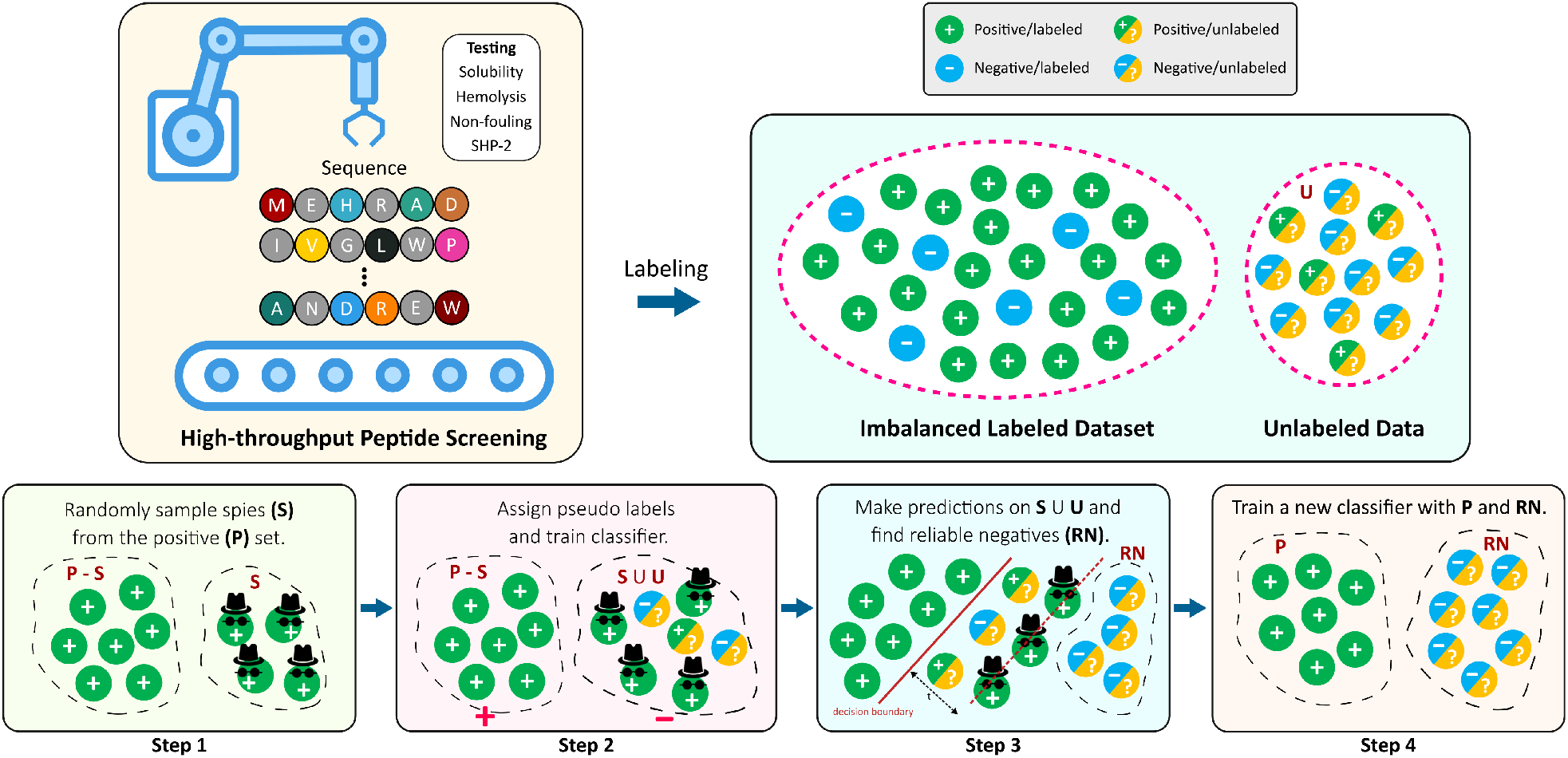
Overview of this work. High-throughput screening methods are commonly good at identifying positive examples, leaving imbalanced datasets (skewed towards the positive class) that are not suitable for supervised learning algorithms. In this work, we use the positive examples only to distinguish between the positive and negative samples using SPY technique.

**Figure 2:**
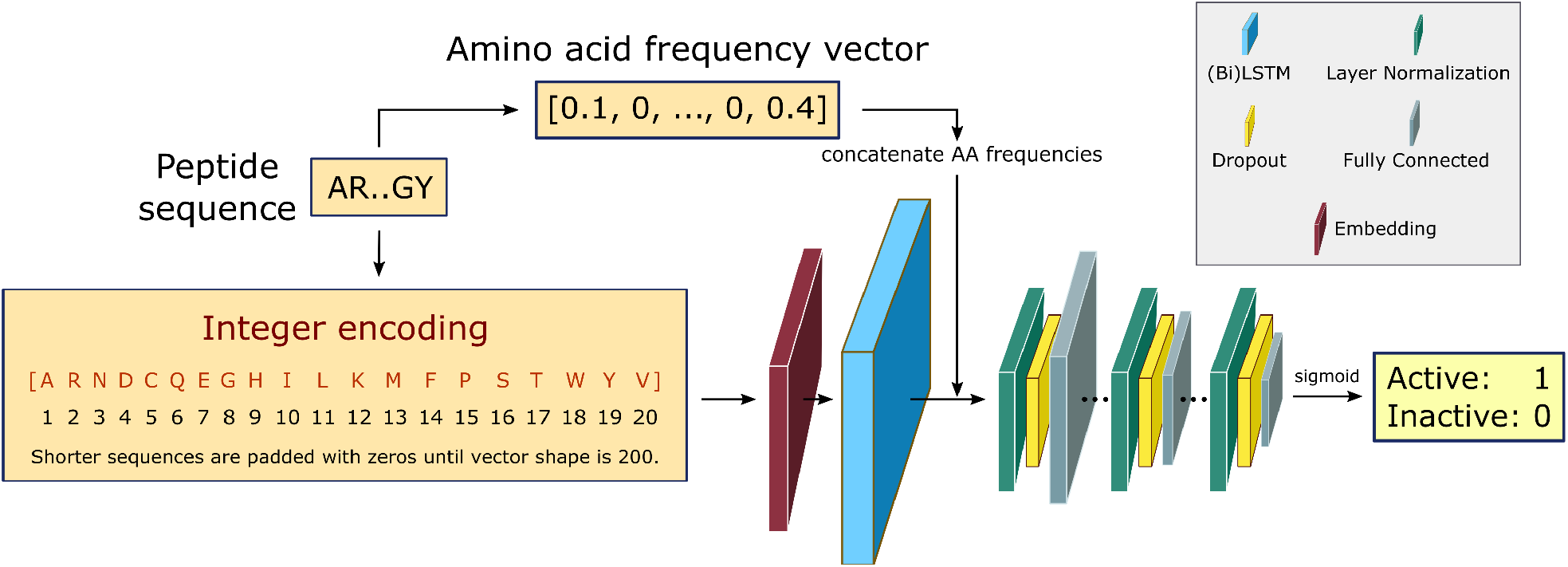
RNN architecture [64]. Padded integer encoded sequences are first fed to a trainable embedding layer, yielding a semantically more compact representation of the input essential amino acids. The use of bidirectional LSTMS and direct inputs of amino acid frequencies prior to the fully connected layers, improves the learning of bidirectional dependency between distant residues within a sequence. The fully connected layers are down-sized in three consecutive steps via layer normalization and dropout regularization. The final layer outputs the probability of being active for the desired training task using a sigmoid activation function.

The input peptide sequences are integer encoded as vectors of shape 200, where the integer at each position in the vector corresponds to the index of the amino acid from the alphabet of the 20 essential amino acids: [A, R, N, D, C, Q, E, G, H, I, L, K, M, F, P, S, T, W, Y, V]. For implementation purposes during the training step, the maximum length of the vector is fixed at 200, padding zeros to shorter length sequences. For those sequences with shorter lengths, zeros are padded to the integer encoding representation to keep the shape fixed at 200 for all examples, to allow input sequences with flexible lengths. Every integer encoded sequence is first fed to an embedding layer with trainable weights, where the indices of discrete symbols (i.e. essential amino acids), into a representation of a fixed-length vector of defined size.

The embedding layer output either goes to a double stacked bi-LSTM layer (for solubility and hemolysis) or a single LSTM layer (for solubility and non-fouling), to identify patterns along a sequence that can be separated by large gaps. The output from the LSTM layer is then concatenated with the relative frequency of each amino acid in the input sequences. This choice is partially based on our earlier work [63], and helps with improving model performance. The concatenated output is then normalized and fed to a dropout layer with a rate of 10%, followed by a dense neural network with ReLU activation function. This is repeated three times, and the final single-node dense layer uses a sigmoid activation function to predict the peptide biological activity as the probability of the label being positive.

The hyperparameters are chosen based on a random search that resulted the best model performance in terms of the area under the receiver operating characteristic curve (AUROC) and accuracy. Readers are encouraged to refer to [64] for more details on the model architecture and its hyperparameters. We compile our Keras model using Adam optimizer [70] with a binary cross-entropy loss function, which is defined as

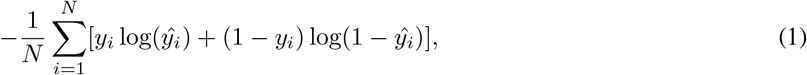

where *y*_*i*_ is the true value of the ith example, *ŷ*_*i*_ is the corresponding prediction, and *N* is the size of the dataset.

### 2.3 Positive-Unlabeled Learning

Let 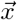 be an example, and *y* ∈ {0, 1} the true binary label for the instance 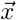. If 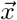 is a positive example, *y* = 1, otherwise *y* = 0. Let *s* = 1, if example 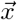 is labeled, and *s* = 0, if 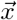 is unlabeled. Only positive examples are labeled (i.e. 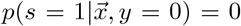). In other words, the probability that a negative example appears in the labeled set is zero. On the other hand, the unlabeled set 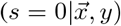 can contain positive 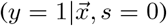 or negative 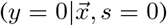 examples. The goal is to learn a probabilistic binary classifier as a function 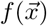, such that 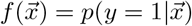, i.e. the conditioned probability of being positive given a feature vector 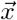.

In this work, we focus on two PU learning strategies; Adapting Base Classifier and Reliable Negative Identification.

#### 2.3.1 Adapting Base Classifier

Adapting base classifier, also known as class prior estimation, are Bayesian-based methods that adapt the base classifier (i.e. SVM) to estimate the expected ratio of positive or negative examples in the unlabeled set. Note that in this work, we use an RNN as our base classifier. This approach simply tries to adjust the probability of being positive estimated by a traditional classifier trained with positive and unlabeled examples, where the unlabeled is treated as the negative class.

The positive likelihood score 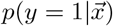 is estimated by Elkan and Noto [71] as

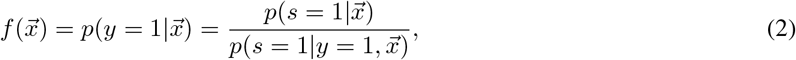

where 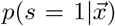 is the likelihood of the example 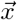 being labeled (thus, being positive), learned from the labeled and unlabeled data. 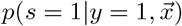 denotes the posterior probability of the example 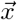, i.e. positive sample being labeled as positive in the training data. Assuming that the labeled positive samples are chosen completely randomly from all positive examples, 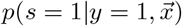 is treated as a constant factor (c) for all the samples, that can be obtained through a validation (held-out) set [53]. This *“selected completely at random”* assumption can be also written as 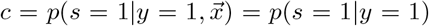, where c is a constant probability that a positive sample is labeled. This assumption is analogous to the *“missing completely at random”* assumption that is made when learning data with missing values [72–74]. Among the empirical estimators for c proposed in [71], we use the following average:

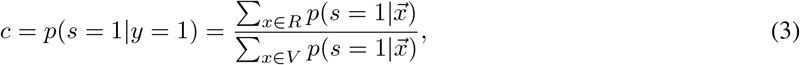

where *V* is the validation set, drawn in the same manner as the training set, and *R* ⊆ *V* is a set of positive examples in *V*. A threshold is adjusted within range (0 − 1/*c*) to discriminate if the sample belongs to the positive or negative class, by maximizing Cohen’s kappa coefficient [75]. It is important to note that the Elkan and Noto [71] algorithm was not developed to handle noisy labeled data. In addition, the theory behind its estimator limits its use to classify conditional distributions with non-overlapping support [76].

#### 2.3.2 Reliable Negative Identification

Reliable negative identification adopts two independent algorithms: 1) identify the reliable negatives (RN) within the unlabeled set given the likelihood and 2) train a binary classifier to distinguish the labeled positive examples from the identified RN set. This approach is based on two assumptions of smoothness and separability, which simply means that all the positive examples are similar to the labeled examples, and that the negative examples are very different from them, respectively [42]. Several techniques have been proposed to extract the reliable negatives or positives from the unlabeled set, such as Spy [43], Cosine-Rocchio [77], Rocchio [44], 1DNF [78], PNLH [79], and Augmented Negatives [80], and DILCA [81].

In this work, we use Spy to find the reliable negatives. First, a small randomly selected group of positive examples (S) are removed and put in the unlabeled data as spies. This allows us to define new datasets *P*_*s*_ and *U*_*s*_, respectively. The percentage of positive instances used as spies is defined by *spy-rate* (in this work, we use 0.2). Then, a classifier *f*_1_ is trained based on *P*_*s*_ and *U*_*s*_. Next, the boundary of RN under the rule that most of the spies are classified as positives is found, based on spy-tolerance (*ϵ*). *ϵ* determines what percentage of spies can remain in the unlabeled set when the decision boundary threshold (*t*_*s*_) is calculated (in this work, we use 0.05). In other words, *t*_*s*_ is the posterior likelihood such that all added spies during training *f*1 are classified as positives. All samples in *U*_*s*_, whose posterior likelihood is smaller than *t*_*s*_ are considered *RN*. Finally, we train a new classifier *f*_2_ given original positive samples (*P*) and the found *RN*.

##### Algorithm 1

Reliable Negative Identification with Spy

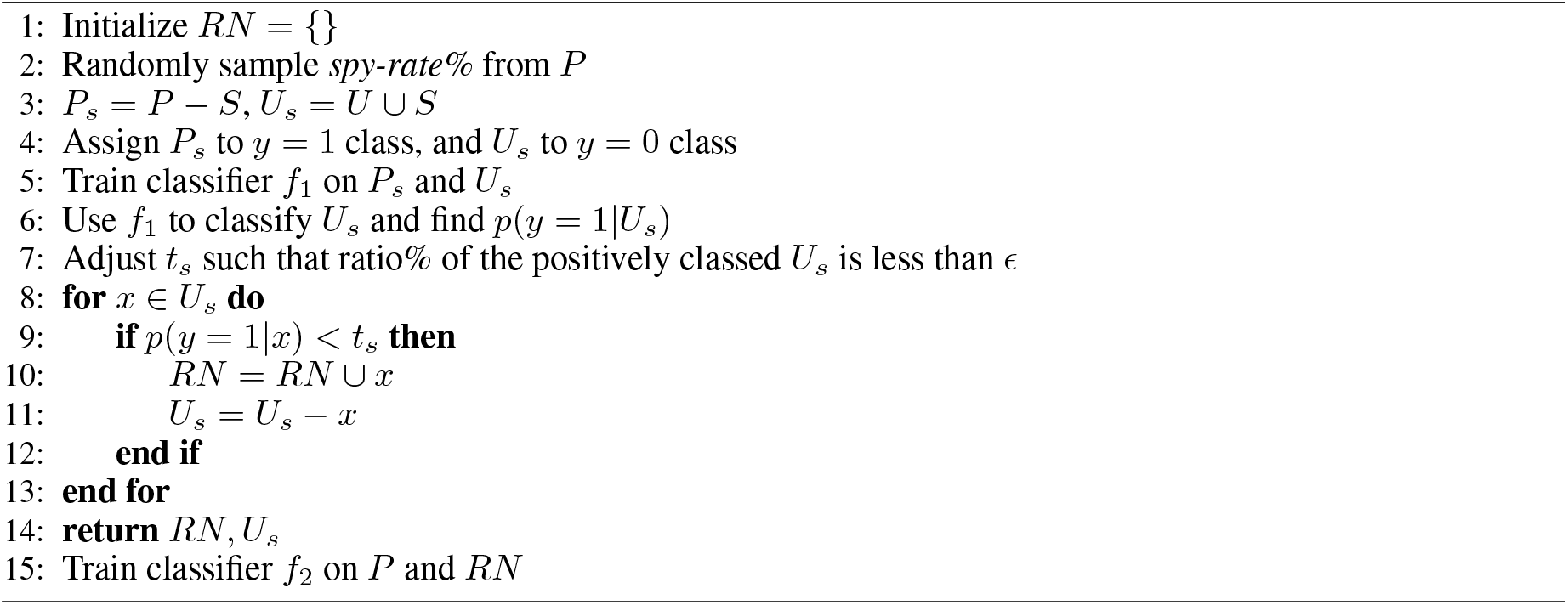

## 3 Results and Discussion

In this section, we evaluate the estimated generalization error of our PU approach, and compare it with the classical PN classification, where both positive and negative examples are available for training. Note that the test data contains unobserved real positive and negative examples. We take two approaches to generate the unlabeled data: 1) Unlabeled Data Generated from Positive and Negatives Samples. In this setting, the unlabeled data is generated from a mixture of known positive and negative examples for each task. 2) Unlabeled Data Generated from Mutated Positive Samples. Given a distribution of positive examples, we generate unlabeled examples by randomly breaking the positive examples into sub-sequences, and filling up a similar-length sequence, with these sub-sequences. Duplicate sequence are removed after the generation step. This allows us to generate the unlabeled data, by creating mutations of the positive examples *without* any knowledge on what the true negative examples are.

### 3.1 Unlabeled Data Generated from Positive and Negatives Samples

Performance comparison between our PU learning methods and classical PN learning for different prediction tasks are presented in Table 2. Results for all the PN models are based on our earlier work in [64]. For every task, we make comparisons of the model accuracy (ACC%), and the area under the receiver operating characteristic curve (AUROC), using the two the Adapting Base classifier, and the Reliable Negative Identification PU methods. Across all prediction tasks, with one exception of Hemolysis and Solubility with the Adapting Base Classifier method, the accuracy of our PU methods are considerably higher than the PN classification. Comparing the two PU methods, it is observed that Reliable Negative Identification outperforms Adapting Base Classifier method for all prediction tasks. Surprisingly, for the non-fouling and SHP-2 predictions, both PU methods outperform the PN classifier.

**Table 1:** Summary of used datasets. For more details, refer to [64].

|  | Hemolysis | Solubility | Non-fouling | SHP-2 |
| --- | --- | --- | --- | --- |
| **Definition** | Hemolysis is the process by which red blood cells (RBCs) rupture and release their contents, mainly Hemoglobin, into the surrounding plasma or extracellular fluid. Based on DBAASP v3 [56]. | Solubility was defined in PROSO II [57] as a sequence that was transfectable, expressible, secretable, separable, and soluble in E. coli system. | Resistance to non-specific interactions. Gathered using the mechanism proposed in [60]. | SHP-2 is a protein encoded by the PTPN11 gene in humans. It is a non-receptor protein tyrosine phosphatase that plays a critical role in various cellular signaling pathways. [62]. |
| **Total size** | 9,316 | 18,453 | 17,185 | 300 |
| **Positive examples** | 19.6% | 47.6% | 20.9% | 50.0% |
| **Length range** | 1-190 AA residues | 19-198 AA residues | 5-198 AA residues | 5 AA residues |

**Table 2:** Performance comparison between PU learning and classical PN learning for different prediction tasks, with the unlabeled data generated from positive and negatives samples. PN models are trained by having access to both positive and negative data, based on our earlier work in [64].

| Task | PU Method | PU |  | PN |  |
| --- | --- | --- | --- | --- | --- |
|  |  | ACC(%) | AUROC | ACC(%) | AUROC |
| Hemolysis | Adapting Base Classifier | 83.1 | 0.78 | 84.0 | 0.84 |
| Hemolysis | Reliable Negative Identification | 84.1 | 0.80 |  |  |
| Non-fouling | Adapting Base Classifier | 93.8 | 0.93 | 82.0 | 0.93 |
| Non-fouling | Reliable Negative Identification | 95.0 | 0.93 |  |  |
| Solubility | Adapting Base Classifier | 53.0 | 0.59 | 70.0 | 0.76 |
| Solubility | Reliable Negative Identification | 86.7 | 0.68 |  |  |
| SHP-2 | Adapting Base Classifier | 84.1 | 0.87 | 83.3 | 0.82 |
| SHP-2 | Reliable Negative Identification | 90.2 | 0.93 |  |  |

### 3.2 Unlabeled Data Generated from Mutated Positive Samples

Table 3 shows performance comparison between our PU learning method and classical PN learning for different prediction tasks. Considering the much better performance of Reliable Negative Identification compared to the Adapting Base Classifier observed in Table 2, we only consider the Reliable Negative Identification PU method for this unlabeled data generation scenario. Note that the solubility model in this setting showed a poor performance, and was excluded in our comparison. Considering the ACC and AUROC reported in Table 3, our PU method is able to reasonably discriminate between the positive and the reliable negatives identified from the generated unlabeled examples.

**Table 3:** Performance comparison between PU learning and classical PN learning for different prediction tasks, with the unlabeled data generated from mutated positive samples. Generated unlabeled is 8 times larger than the positive size. PN models are trained by having access to both positive and negative data, based on our earlier work in [64].

| Task | PU Method | PU |  | PN |  |
| --- | --- | --- | --- | --- | --- |
|  |  | ACC(%) | AUROC | ACC(%) | AUROC |
| Hemolysis | Reliable Negative Identification | 76.8 | 0.75 | 84.0 | 0.84 |
| Non-fouling | Reliable Negative Identification | 94.1 | 0.87 | 82.0 | 0.93 |
| SHP-2 | Reliable Negative Identification | 84.8 | 0.91 | 83.3 | 0.82 |

It is important to note that with the unlabeled data generation, we can control how large the size of the generated unlabeled examples are. The generated unlabeled:labeled ratio reported in Table 3 is fixed at 8.0. Next, we investigate the effect of unlabeled:labeled ratio on the performance of Reliable Negative Identification strategy across all prediction tasks in Figure 3. Each point represents the average value of AUROC and ACC% (left and right panel, respectively) over 6 models trained with a different choice of randomly selected spy positives, and error bars show the magnitude of the standard deviation. Horizontal dashed lines show the performance of the PN classifier for each task represented as a baseline for performance comparison. With very small generated unlabeled samples (i.e. unlabeled:labeled ratio *≈* 2.0), the exploration of new examples that can qualify as reliable negatives will be largely limited. Thus, the trained *f*_2_ classifier has a significantly lower performance compared to the baseline PN classifier and to the other PU models trained with higher generated unlabeled:labeled ratios. With larger unlabeled:labeled ratios (i.e. > 10.0), we see a better prediction performance across all the tasks. There are two significant observations; 1. With more unlabeled sequences generated, the trained PU models have a competitive performance with the PN models. In specific, for binding against SHP2, we observe that the PU model beats the PN classifier in both AUROC and ACC%. 2. Surprisingly, the PU models become more confident in their predictions with the increase of the unlabeled:labeled ratio (compare magnitude of error bars in Figure 3). This can bring a major advantage in implementing our approach in a generative setting, where we can predict the properties of new peptide sequences without having to worry much about the class imbalance between the positive and the negative examples, which can majorly reduce model performance, if the learning is supervised.

**Figure 3:**
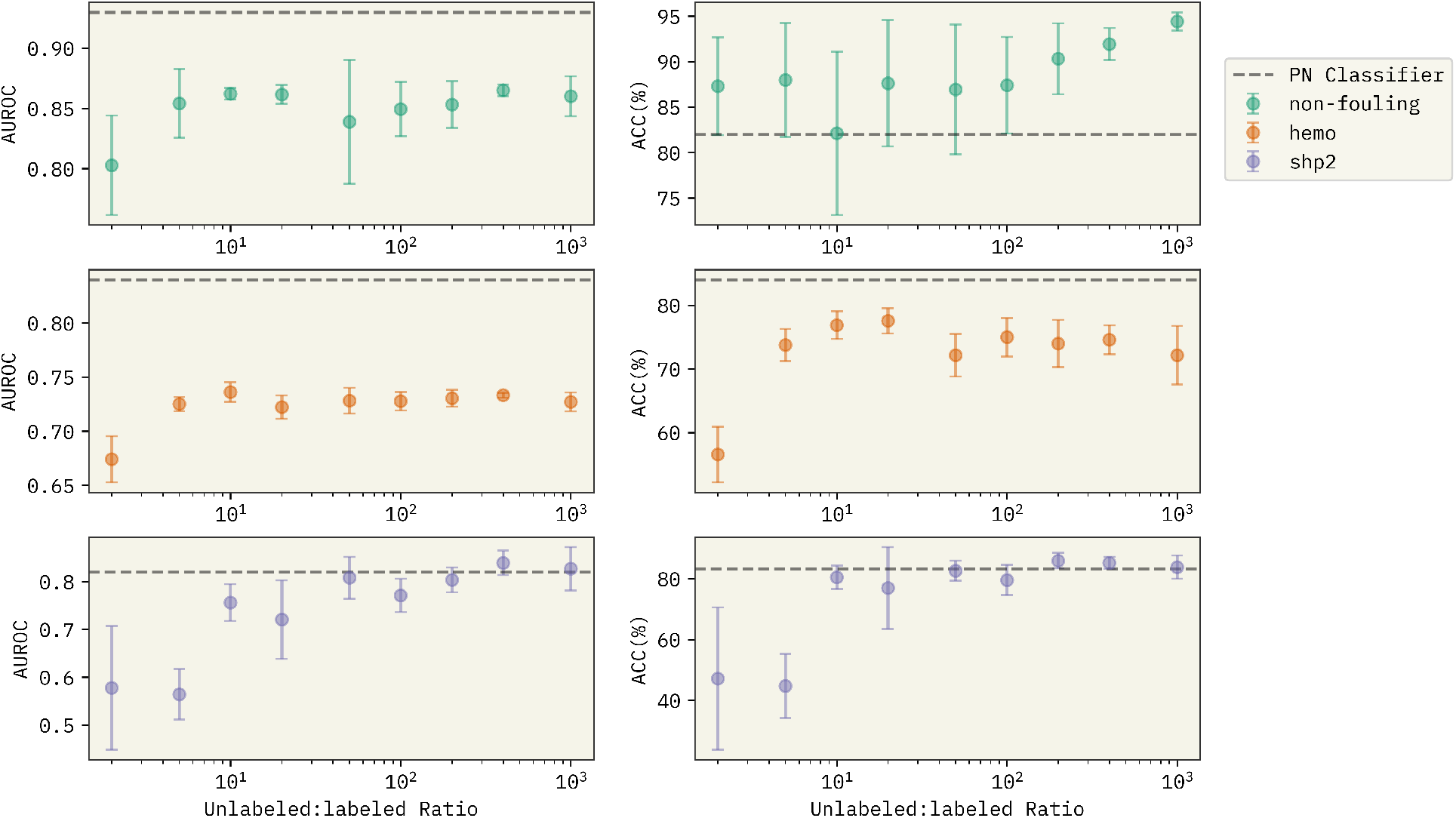
Effect of generated unlabeled:labeled ratio on the performance of the Reliable Negative Identification strategy for the three prediction tasks. Horizontal dashed lines show the performance of the PN classifier from Table 3 used as a baseline for comparison. At the low ratio regime, the pool of unlabeled data is not big enough to obtain promising candidates as reliable negatives. With larger unlabeled:labeled ratios, the PU model gets to identify a better choice of sequences as reliable negatives, despite the major existing class imbalance in the traning data.

Comparing AUROC and ACC in Tables 2 and 3, we observe that Reliable Negative Identification with mutated positive samples has a relative lower performance compared to the other scenario, where the unlabeled data is generated from a distribution of positive and negative examples. Despite this minor lower performance, using the new unlabeled sequence generation, one can explore the newly unlabeled samples, and make predictions on peptide properties by only having access to the examples from one class (i.e. positive). The sequence-based peptide property prediction in this work is limited to four different tasks. However, with the positive data available, this work can be further extended to developing predictive models for inferring other peptide properties.

## 4 Conclusions

We’ve showed a semi-supervised learning framework to infer the mapping from peptides’ sequence to function for properties such as hemolysis, solubility, non-fouling, and binding against SHP-2. Our positive unlabeled learning method aims at identifying likely negative candidates(reliable negatives), from the generated unlabeled sequences, given random permutations of subsequences within the available positive samples. The reliable negative identification strategy is agnostic with respect to the model architecture used, giving generality. Our method will be most beneficial in biology screening experiments, where most high-throughput screening methods solely focus on identifying the positive example. All PU models showed a comparative predictive ability and robustness across the different prediction tasks, when compared to training with both positive and negative examples. This learning strategy can provide a robust feasible path towards estimating how amino acids positional substitutions can affect peptide’s functional response for unknown sequences, and accelerate the design and discovery of novel therapeutics.

## Acknowledgements

Research reported in this work was supported by the National Institute of General Medical Sciences of the National Institutes of Health under award number R35GM137966. We thank the Center for Integrated Research Computing (CIRC) at University of Rochester for providing computational resources and technical support.

## Data and Code Availability

All data and code used to produce results in this study are publically available in the following GitHub repository: https://github.com/ur-whitelab/pu-peptides.

